# Flow Cytometry Strategies for Rapidly Characterizing Heterogeneous Adipocyte Populations in 3D In Vitro Constructs

**DOI:** 10.1101/2024.01.30.578065

**Authors:** Michael Struss, Golnaz Anvari, Evangelia Bellas

## Abstract

Adipose tissue (AT) is an endocrine organ that regulates whole body metabolism and supports energy needs of other tissues. Two key adipose tissue functions are insulin-stimulated glucose uptake and lipid metabolism. As the prevalence of metabolic diseases, such as obesity, continue to rise, there is a growing need for new methods to study adipose tissue and its main cell type, adipocytes. Adipocytes are unique cells, distinguished by their large spherical shape housing large lipid droplet(s). For many in vitro models (and in tissues), adipocytes are derived from a heterogenous population of precursor cells, leading to varying degrees of adipogenesis and adipocyte maturation. Characterization of such populations can be challenging because often the average result does not account for the complexity of the various sub-populations. Common single cell characterization methods provide data based gene and protein expression but do not account for the morphological variability in cells such as adipocytes at different stages of maturation, and are expensive to run. More traditional methods, such as microscopy or colorimetric assays, are often time consuming with intrinsic challenges due to overlapping or coincident features or lose single cell adipocyte details due to the destructive nature of the assay. Here, we show how flow cytometry can be used to characterize adipocyte populations while preserving critical details at the individual adipocyte level.

- This protocol provides multiple workflows for indirect measurements of lipogenesis (lipid accumulation), protein content (branched actin formation), and adipocyte functions (insulin-stimulated glucose uptake).
- The flow cytometry workflows presented in this work show the effectiveness of binning individual adipocytes based on their level of maturity and allow for comparisons within subpopulations traditional methods cannot provide.

**Graphical abstract:** 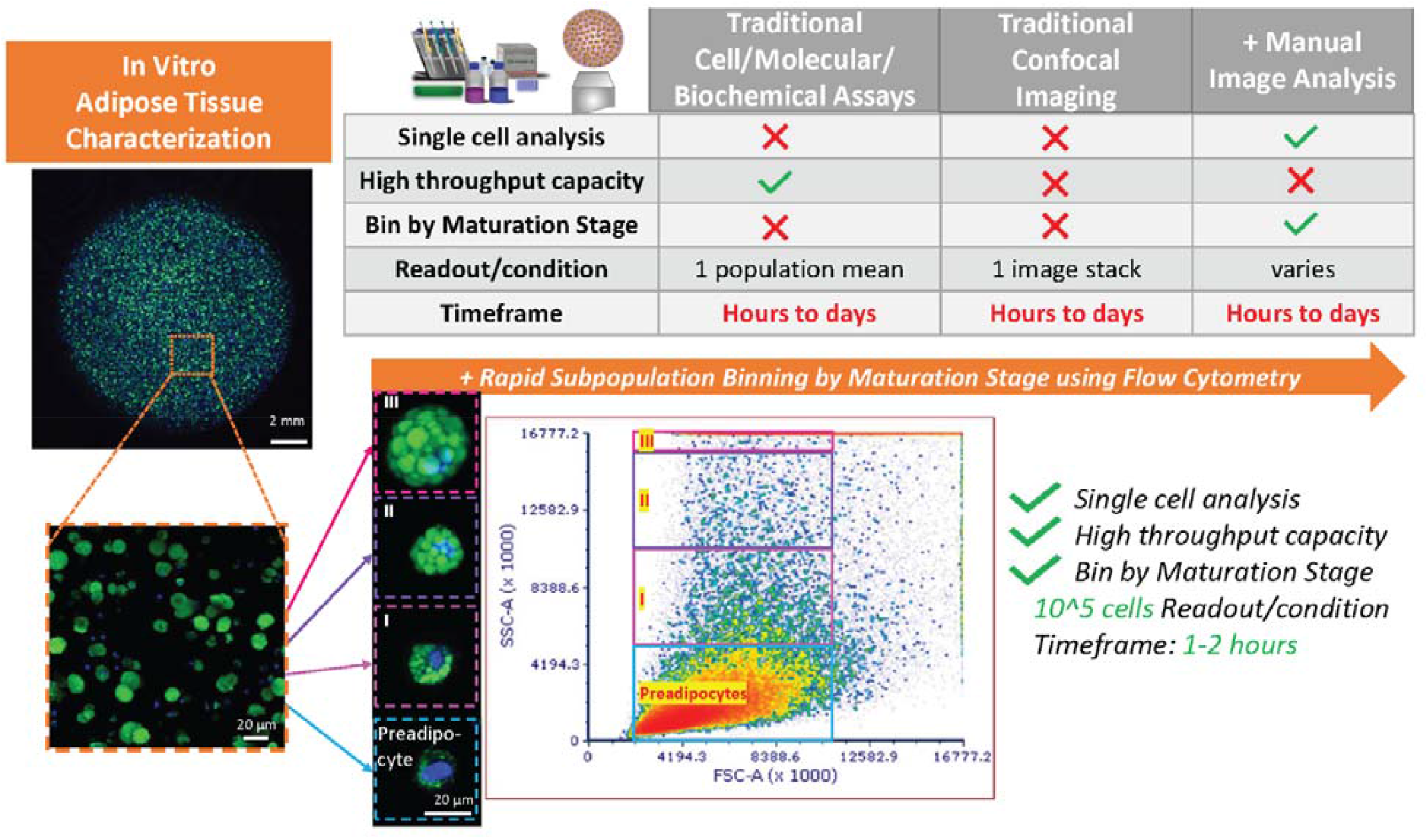

|  |  |
| --- | --- |
| Subject area | Engineering |
| More specific subject area | Tissue Engineering |
| Name of your method | Flow Cytometry Strategies for Binning Adipocyte Subpopulations |
| Name and reference of original method | n/a |
| Resource availability | n/a |

## Method details

### Introduction

Obesity, broadly defined as excess adipose tissue, is a growing epidemic, where the prevalence in adults in the United States is 42.4% in 2017-2018[1]. As a result of this growing epidemic, researchers are developing new approaches to understanding adipose tissue and the main cell in adipose tissue, adipocytes. Adipocytes are unique cells, with a large spherical shape to accommodate lipid storage. Therefore, for in vitro studies, three-dimensional (3D) environments are needed to support the growth into large, round, lipid-containing adipocytes[2] and provide a more accurate model for examining the interactions between adipocytes and their environment[3–13]. Lipogenesis, the accumulation, and storage of intracellular lipid droplets within adipocytes, is a hallmark of adipocytes. Intracellular lipids are stained with lipophilic dyes such as Oil Red O, Fluorinated Boron-Dipyrromethene 493/503 (BODIPY 493/503), or Nile Red, and visualized by light or fluorescent microscopy. These dyes (Oil Red O, Nile Red) can be eluted and quantified by colorimetric assays[14–17]. During early adipogenesis, actin stress fibers are disrupted to allow for this lipid accumulation, and actin-related protein 3 (ARP3) mediated branched cortical actin formation facilitates further lipid accumulation[18]. Actin branching is necessary for proper cortical actin formation and plays an essential role in intracellular vesicle translocation to the cell membrane, such as glucose transporter type 4 (GLUT4)[19]. Therefore, adipocyte maturation involves an accumulation of intracellular lipids and branched cortical actin proteins essential for adipocyte function. However, adipogenic maturation is an intrinsically heterogeneous process making it difficult to compare individual adipocytes to the population using microscopy or colorimetric assays alone[20,21].

High magnification and confocal microscopy give precise details about intracellular lipid droplet morphology and organization within adipocytes. However, this 3D imaging can be very time-consuming depending on the thickness of the sample and the number of fluorescent channels imaged. Further, image quantification of these intracellular features is inherently challenging to distinguish overlapping or coincident lipid droplet features. Conversely, colorimetric or biochemical assays provide quick assessments of adipocyte populations within 3D adipose tissue (AT) constructs. However, single-cell adipocyte details are lost from the destructive nature of the assay.

The limitations of using microscopy and colorimetric assays to capture single cell details are not restricted to measuring adipocyte lipid droplet content. Similar issues arise when quantifying other intracellular features either visually by immunofluorescence or western blot. These same issues also apply to functional assays such as glucose uptake by visualizing uptake of a fluorescent glucose analog or biochemically by sampling media for depleted glucose. Therefore, there is a need for improved, inexpensive methods to quantify 3D adipocyte populations at the single-cell level to obtain information about the heterogenous population.

In this work, flow cytometry was used to characterize adipocyte populations while preserving critical details at the individual adipocyte level. Here, we present how flow cytometry enables the indirect measurements of lipogenesis (lipid accumulation), protein content (branched actin formation), and adipocyte functions (insulin-stimulated glucose uptake). By binning adipocytes into subpopulations based on side scatter (SSC), comparisons between subpopulations can be made to better understand how adipocyte morphology and function change based on maturity. The 3D adipocyte flow cytometry workflows demonstrate the utility of characterizing individual adipocytes based on their maturity level, allowing for comparisons not possible using traditional methods, such as microscopy or colorimetric assays, alone.

## Method details

### Gelatin Methacryloyl (GelMA) synthesis

10 grams of gelatin (Sigma-Aldrich, type A, 300 bloom) were dissolved in 100 mL of 0.25 M sodium bicarbonate buffer (Sigma-Aldrich) in a 500 mL glass bottle at 50°C on a stirring hot plate until completely dissolved. Once dissolved, 2.5 mL (2.5% v/v) methacrylic anhydride (Sigma-Aldrich, 99%) was added dropwise at a flow rate of 0.6 mL/minute to the gelatin solution and allowed to react for 1 hour. During the reaction, the pH was adjusted to 9 using 1M NaOH (Fisher) and protected from light.

To ensure reproducibility, the methacrylation reaction should be maintained at 50°C and pH of 9.

We used a digital temperature probe and pH probe.

The gelatin methacryloyl (GelMA) solution was diluted at 1:4 in prewarmed 1X phosphate buffer saline (PBS, Fisher Scientific), and placed in dialysis tubes (Sigma-Aldrich, 12-14 kDa MWCO). The GelMA solution was dialyzed for 3 days against 3 L of 1X PBS followed by 3 days against 3 L of deionized water. All dialysis was performed at 55°C on a stirring hot plate and protected from light. After dialysis, the GelMA solution was sterile filtered through a 0.22 µm filter, aliquoted into sterile tubes, snap-frozen, and lyophilized. The lyophilized GelMA was stored at -80°C until needed.

### Two-dimensional (2D) human mesenchymal stem cell (hMSC) culture and adipogenic induction

Human bone marrow-derived mesenchymal stem cells (hMSCs) were isolated from healthy adult male bone marrow, purchased from Lonza Walkersville Inc. (Walkersville, MD, Cat# 1M-105). Cells were maintained at 37°C and 5% CO_2_ for all experiments. HMSCs were expanded until passage 4 in low glucose media containing 10% v/v fetal bovine serum (FBS, Sigma-Aldrich) and 1% v/v penicillin/streptomycin (P/S, HyClone). When hMSCs were confluent at passage 4, growth media was exchanged for adipogenic induction media (Table 1) and cultured for 7 days.

**Table 1:** Adipogenic induction media components.

| **Table 1: Adipogenic induction media components** |  |  |  |  |  |  |  |
| --- | --- | --- | --- | --- | --- | --- | --- |
| **Name** | **Vendor** | **Catalog Number** | **Molecular Weight** | **Stock Solution** | **Stock Solution Preparation** | **Final Concentration in Media** | **Per 100 mL of media** |
| Dulbecco's Modified Eagle Medium/Nutrient Mixture F-12 | Corning | MT10092 CM | - | 1X | - | - | - |
| Fetal Bovine Serum (FBS) | Sigma-Aldrich | 12306C | - | - | - | 3% v/v | 3 mL |
| Penicillin/Streptomycin (P/S) | HyClone | SV30010 | - | 100X | - | 1% v/v | 1 mL |
| Rosiglitazone | Cayman Chemicals | 71740 | 357.43 | 10 mM | 10 mg in 2.8 mL DMSO | 2 µM | 20 µL |
| Isobutylmethylxanthine | Thermo Scientific | I7018 | 222.24 | 50 mM | 250 mg in 19.4 mL H<sub>2</sub>O+ 2.15 mL 1M NaOH | 500 µM | 1000 µL |
| Dexamethasone | Acros Organics | 50022 | 392.46 | 0.5 mM | 10 mg in 52 ml EtOH | 1 µM | 200 µL |
| D-calcium pantothenate | Acros Organics | 2433011 000 | 238.27 | 84 mM | 100 mg in 5 ml H<sub>2</sub>O | 17 µM | 20 µL |
| D-Biotin | AlfaAesar | 58855 | 65 | 65 mM | 160 mg in 10 ml (9 ml H<sub>2</sub>O+1 ml 1M NaOH) | 33 µM | 55 µL |
| Insulin | Sigma-Aldrich | I9278 | 1.7 | 1.7 mM | 10 mg/ml | 20 nM | 50 µL |

### Three-dimensional (3D) adipocyte encapsulation and tissue culture

After 7 days of adipogenic induction, adipocytes were washed with 1X PBS, detached via trypsin (Gibco), and resuspended at 8 million cells/mL in a 5% w/v gelatin methacryloyl (GelMA) hydrogel solution containing 0.05% w/v photoinitiator, lithium phenyl-2,4,6-trimethylbenzoylphosphinate (LAP), described in Table 2.

**Table 2:**
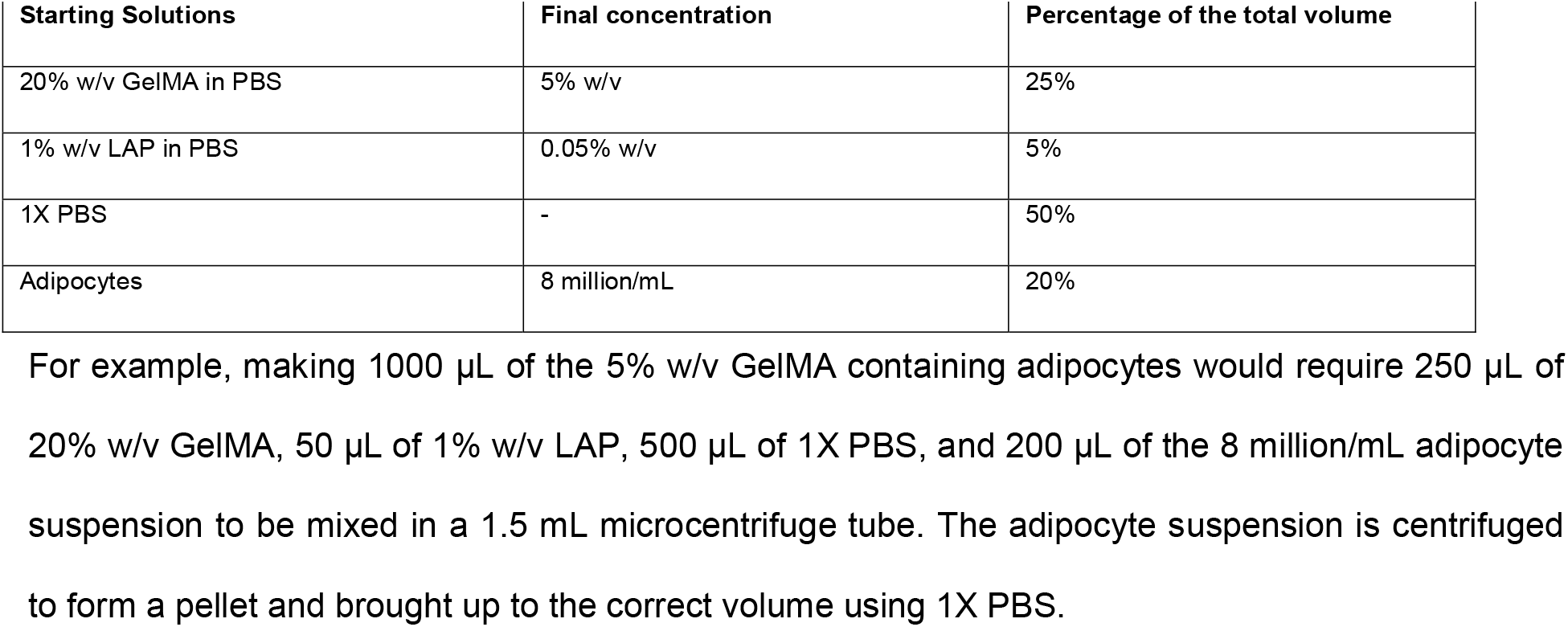
5% w/v gelatin methacryloyl (GelMA) formulation.

For example, making 1000 µL of the 5% w/v GelMA containing adipocytes would require 250 µL of 20% w/v GelMA, 50 µL of 1% w/v LAP, 500 µL of 1X PBS, and 200 µL of the 8 million/mL adipocyte suspension to be mixed in a 1.5 mL microcentrifuge tube. The adipocyte suspension is centrifuged to form a pellet and brought up to the correct volume using 1X PBS.

The adipocyte containing hydrogel precursor solution (30 µL volume) was pipetted into spherical injection molds and photocrosslinked with 405 nm light (15 mW) for 2 minutes. The adipocyte containing GelMA spherical hydrogels (referred to now as *AT constructs*) were transferred to T25 cell culture flasks. Media was replenished every other day and AT constructs were maintained for up to 28 days.

### Fluorescent staining for flow cytometry

Six AT constructs were fixed in 1.5 mL of 4% (v/v) paraformaldehyde (Thermo Scientific) in 35mm dishes for 1 hour and washed 3 times with 3 mL of 1X PBS. After fixation, 2 AT constructs were moved to 1.5 mL microcentrifuge tubes and stained with 200 µL of 5 µg/mL BODIPY 493/503 (1:200 dilution) and 5 mM Deep Red Anthraquinone 5 (DRAQ5, 1:1000 dilution) in 1X PBS for 1 hour. After staining, AT constructs were washed 3 times with 200 µL 1X PBS and digested in 100 µL of 35 mg/mL collagenase type I (Gibco) in 1X PBS for 30 minutes at 37°C. The digested AT construct is homogenized by pipetting up and down thoroughly until no visible chunks can be seen. After homogenization, the adipocyte suspension is brought to a final volume of 300 µL using 1X PBS and transferred to a 5 mL round-bottom tube (Falcon). More details about fluorescent stains can be found in Table 3.

**Table 3:** Staining reagent formulations.

| Target | Reagent | Excitation/<br>Emission | Stock<br>Solution | Working<br>Solution | Manufacturer/<br>Catalog Number | Antibody RRID |
| --- | --- | --- | --- | --- | --- | --- |
| Nucleus | DRAQ5 | 647/681 | 5 mM | 5 μM<br>(1:1000) in<br>1X PBS | Thermo Scientific/62254 | - |
| Nucleus | Hoechst<br>33342 | 361/497 | 20 mM | 20 μM<br>(1:1000) in<br>1X PBS | Thermo Scientific/62249 | - |
| Lipids | BODIPY<br>493/503 | 493/503 | 1 mg/mL in<br>EtOH | 5 ug/mL<br>(1:200) in<br>1X PBS | Invitrogen/D3922 | - |
| Glucose | 2-NBDG | 465/540 | 10 mM in<br>DMSO | 10 μM<br>(1:1000) in<br>1X PBS | Invitrogen/N13195 | - |
| Actin related<br>protein 3<br>(ARP3) | Rabbit anti-Human<br>ARP3 | - | - | (1:100) in<br>1X PBS | Cell Signaling/4738 | AB_2221973 |
| Rabbit IgG | Goat anti-Rabbit,<br>Alexa<br>Fluor™ 488 | 498/520 | 2 mg/mL | 2 ug/mL<br>(1:1000) in<br>1X PBS | Invitrogen/A11008 | AB_143165 |
| Rabbit IgG | Goat anti-Rabbit,<br>Alexa<br>Fluor™ 546 | 556/573 | 2 mg/mL | 2 ug/mL<br>(1:1000) in<br>1X PBS | Invitrogen/ A11010 | AB_2534077 |
| Glucose<br>transporter 4<br>(GLUT4) | Mouse anti-Human<br>GLUT4 | - | - | (1:100) in<br>1X PBS | Cell Signaling/2213 | AB_823508 |
| Mouse IgG | Goat anti-Mouse,<br>Alexa<br>Fluor™ 594 | 590/617 | 2 mg/mL | 2 ug/mL<br>(1:1000) in<br>1X PBS | Invitrogen/A11005 | AB_2534073 |

### Fluorescent staining for confocal imaging

Six AT constructs were fixed in 1.5 mL of 4% (v/v) paraformaldehyde (Thermo Scientific) in 35mm dishes for 1 hour and washed 3 times with 3 mL of 1X PBS. After fixation, individual AT constructs were moved to mL microcentrifuge tubes and stained with 100 µL of 5 µg/mL BODIPY 493/503 (1:200 dilution) and 20 mM Hoechst 33342 (1:1000 dilution) in 1X PBS for 1 hour. After staining, AT constructs were washed 3 times with 100 µL 1X PBS and stored in 1X PBS at RT until imaged. More details about fluorescent stains can be found in Table 3.

### Immunofluorescent (IF) staining for flow cytometry

Six AT constructs were fixed with 1.5 mL of 4% (v/v) paraformaldehyde (Thermo Scientific) in 35mm dishes for 3 hours, washed 3 times with 3 mL of 1X PBS, and permeabilized for 1 hour using 1.5 mL of 1% v/v Triton X (Sigma-Aldrich).

We found that fixing AT constructs for 3 hours improved AT construct stability during the IF staining. After fixation and permeabilization, 2 AT constructs were moved to 1.5 mL microcentrifuge tubes and blocked with 200 µL of 3% bovine serum albumin (BSA, Fisher Scientific) in 1X PBS overnight. Next, the AT constructs were incubated overnight with 200 µL of rabbit anti-human actin-related protein 3 (ARP3, 1:100 dilution, RRID: AB_2221973) primary antibody in 3% BSA in 1X PBS overnight. After primary antibody incubation, AT constructs were washed 3 times with 200 µL of 1X PBS and incubated in 200 µL 1X PBS overnight. After washing, the 2 AT constructs were incubated with 200 µL of goat anti-rabbit, Alexa Fluor™ 488 (1:1000 dilution, RRID: AB_143165) secondary antibody in 3% BSA in 1X PBS overnight. After secondary antibody incubation, the AT constructs were washed 3 times with 200 µL of 1X PBS and stained with 200 µL of 5 mM DRAQ5 (1:1000 dilution) in 1X PBS. After nuclear staining, AT constructs were washed 3 times with 200 µL 1X PBS and digested in 100 µL of 35 mg/mL collagenase type I (Gibco) in 1X PBS for 30 minutes at 37°C. The digested AT construct is homogenized by pipetting up and down thoroughly until no visible chunks can be seen. After homogenization, the adipocyte suspension is brought to a final volume of 300 µL using 1X PBS and transferred to a 5 mL round-bottom tube (Falcon). More details about IF reagents can be found in Table 3.

### Immunofluorescent (IF) staining for confocal imaging

Six AT constructs were previously fixed with 1.5 mL of 4% (v/v) paraformaldehyde (Thermo Scientific) in 35mm dishes for 3 hours, washed 3 times with 3 mL of 1X PBS, and permeabilized for 1 hour using 1.5 mL of 1% v/v Triton X (Sigma-Aldrich).

We found that fixing AT constructs for 3 hours improved AT construct stability during the IF staining. After fixation and permeabilization, individual AT constructs were moved to 1.5 mL microcentrifuge tubes and blocked with 200 µL of 3% bovine serum albumin (BSA, Fisher Scientific) in 1X PBS overnight. Next, individual AT constructs were incubated overnight with 100 µL of rabbit anti-human actin-related protein 3 (ARP3, 1:100 dilution, RRID: AB_2221973) primary antibody or 100 µL of mouse anti-human glucose transporter 4 (GLUT4, 1:100 dilution, RRID: AB_823508) primary antibody in 3% BSA in 1X PBS overnight. After primary antibody incubation, individual AT constructs were washed 3 times with 100 µL of 1X PBS and incubated in 100 µL 1X PBS overnight. After washing, individual AT constructs were incubated with 100 µL of goat anti-rabbit, Alexa Fluor™ 594 (1:1000 dilution, RRID: AB_2534077) secondary antibody or goat anti-mouse, Alexa Fluor™ 594 (1:1000 dilution, RRID: AB_2534073) secondary antibody in 3% BSA in 1X PBS overnight. After secondary antibody incubation, individual AT constructs were washed 3 times with 100 µL of 1X PBS and stained with 100 µL of 20 mM Hoechst 33342 (1:1000 dilution) and 5 µg/mL BODIPY 493/503 (1:200 dilution) in 1X PBS.

Hoechst 33342 (361 nm excitation/497 nm emission) was used instead of DRAQ5 (647 nm excitation/681 nm emission) for confocal imaging to avoid overlapping signal from Alexa Fluor™ 546 and 594 (556 nm excitation/573 nm emission; 590 nm excitation/617 nm emission).

After fluorescent staining, AT constructs were washed 3 times with 100 µL 1X PBS and stored in 1X PBS at RT until imaged. More details about fluorescent stains can be found in Table 3.

**Table 4:** Troubleshooting adipocyte flow cytometry.

| Problem | Solution | Reference |
| --- | --- | --- |
| Mature buoyant adipocytes are not collected | - Collection probe stirring ON<br>- Sample agitation ON<br>- Phase separate by centrifugation or buoyancy, separate the buoyant upper layer of mature adipocytes, run flow cytometry on either layer | - [27]<br>- [27], [28]<br>- [27], [28] |
| Adipocyte rupture | - Slow fluidics (6 μL/min is the lowest flow rate for this flow cytometer)<br>- Increase probe nozzle size<br>- Gently pipette up and down when digesting AT construct<br>- Run flow cytometry samples at room temperature | - [27]<br>- [27]<br>- n/a<br>- [27] |
| Excluding debris | - Nuclear staining (Fig. 1)<br>- Running flow cytometry on 2D samples (Supplementary Fig. 1) | - n/a<br>- n/a |
| Gating adipocyte clumps/doublets | - Using signal width (Fig. 1)<br>- Using FSC-A vs. FCS-H<br>- Using SSC-H vs. SSC-W | - n/a<br>- [27], [28], [24]<br>- [28] |
| Adipocytes clogging collection probe | - Longer enzymatic digestion time at 37°C<br>- Extensive pipetting to break up clumps<br>- Using SSC-A vs. time to monitor if a clog has occurred | - n/a<br>- n/a<br>- n/a |
| Removing debris before running flow | - Centrifugation and resuspension with fresh buffer<br>- Cell strainer | - [28], [24]<br>- [27] |
| Hydrogel does not digest | - Depending on the type of hydrogel/biomaterial, some may not be able to be fixed prior to enzymatic digestion (collagen hydrogels in our previous experience)<br>- Increasing digestion enzyme concentration<br>- Longer enzymatic digestion time at 37°C | - n/a<br>- n/a<br>- n/a |

### Insulin-stimulated glucose uptake assay for flow cytometry

AT constructs were first insulin starved for 6 hours in adipogenic induction media without insulin. After 6 hours, media was replaced with adipogenic induction media containing insulin and 20 µM fluorescent analog of glucose, 2-(*N*-(7-Nitrobenz-2-oxa-1,3-diazol-4-yl)Amino)-2-Deoxyglucose (2-NBDG, Invitrogen) and incubated for 30 minutes. After 30 minutes, AT constructs were placed in 35 mm dishes, washed 3 times with 3 mL of 1X PBS, and fixed with 1.5 mL of 4% (v/v) formaldehyde for 1 hour at RT. After fixation, 2 AT constructs were moved to 1.5 mL microcentrifuge tubes and stained with 200 µL of 20 mM DRAQ5

(1:1000 dilution) in 1X PBS for 1 hour. After staining, AT constructs were washed 3 times with 200 µL 1X PBS and digested in 100 µL of 35 mg/mL collagenase type I (Gibco) in 1X PBS for 30 minutes at 37°C. The digested AT construct is homogenized by pipetting up and down thoroughly until no visible chunks can be seen. After homogenization, the adipocyte suspension is brought to a final volume of 300 µL using 1X PBS and transferred to a 5 mL round-bottom tube (Falcon). More details about 2-NBDG can be found in Table 3.

### Confocal microscopy and image processing

Images were acquired with a laser scanning confocal mode of the hybrid confocal, multiphoton microscope (Olympus FluoView FV1200) using a 30X (UPLSAPO30XSIR, 1.05 NA, Olympus) oil immersion inverted objectives with four laser units (405, 488, 543, and 635 nm.), and four photomultipliers (PMT) detectors. Z-stacks (thickness ∼ 50 µm, step-size: 5 µm, 12.5 µs/pixel, 1024x1024) were captured at least 20 µm above the glass coverslip to ensure visualized cells were in a 3D environment and not in contact with the glass coverslip. When appropriate, brightness and contrast were applied equally to all conditions to improve signal and reduce background. FIJI was used for all image processing [22]. Individual adipocytes were isolated from confocal image z-stacks. For BODIPY 493/503 fluorescent images, z-stack slices were z-projected using the Max Intensity method (Fig. 3b, d). For ARP3 and GLUT4 immunofluorescence images were displayed z-stacks in Fig. 4b, 5b, and as individual slices in Fig. 4d, 5d.

### Flow cytometry instrument setup

Flow cytometry was performed using an Accuri C6 flow cytometer (BD Biosciences) with a 24-tube well plate configuration. Data were acquired using BD CSampler analysis software (BD Biosciences). The flow cytometer contains 3 blue (488 nm) and 1 red (640 nm) laser configuration with 4 filter/detector sets (FL1 – 533/30, FL2 – 585/40, FL3 – 670 LP, FL4-675/25). Nuclear staining (DRAQ5) was detected using the FL4-A channel, while ARP3 immunofluorescence and insulin-stimulated glucose uptake fluorescence were detected using the FL1-A channel. BODIPY 493/503 fluorescence was detected in the FL2-A channel.

The flow cytometer limits were set to collect 300 µL of the sample. A vertical gate was placed at 10^3 in the FL4-A channel histogram, and the machine was set not to collect any events with a fluorescent value less than 10^3 in the FL4-A channel to minimize events from debris. Lastly, the flow cytometer fluidics were set to the lowest possible flow rate of 6 µL/min to avoid adipocyte lysis. More details about the gating strategy can be found in the representative results section (Fig. 1).

**Figure 1.**
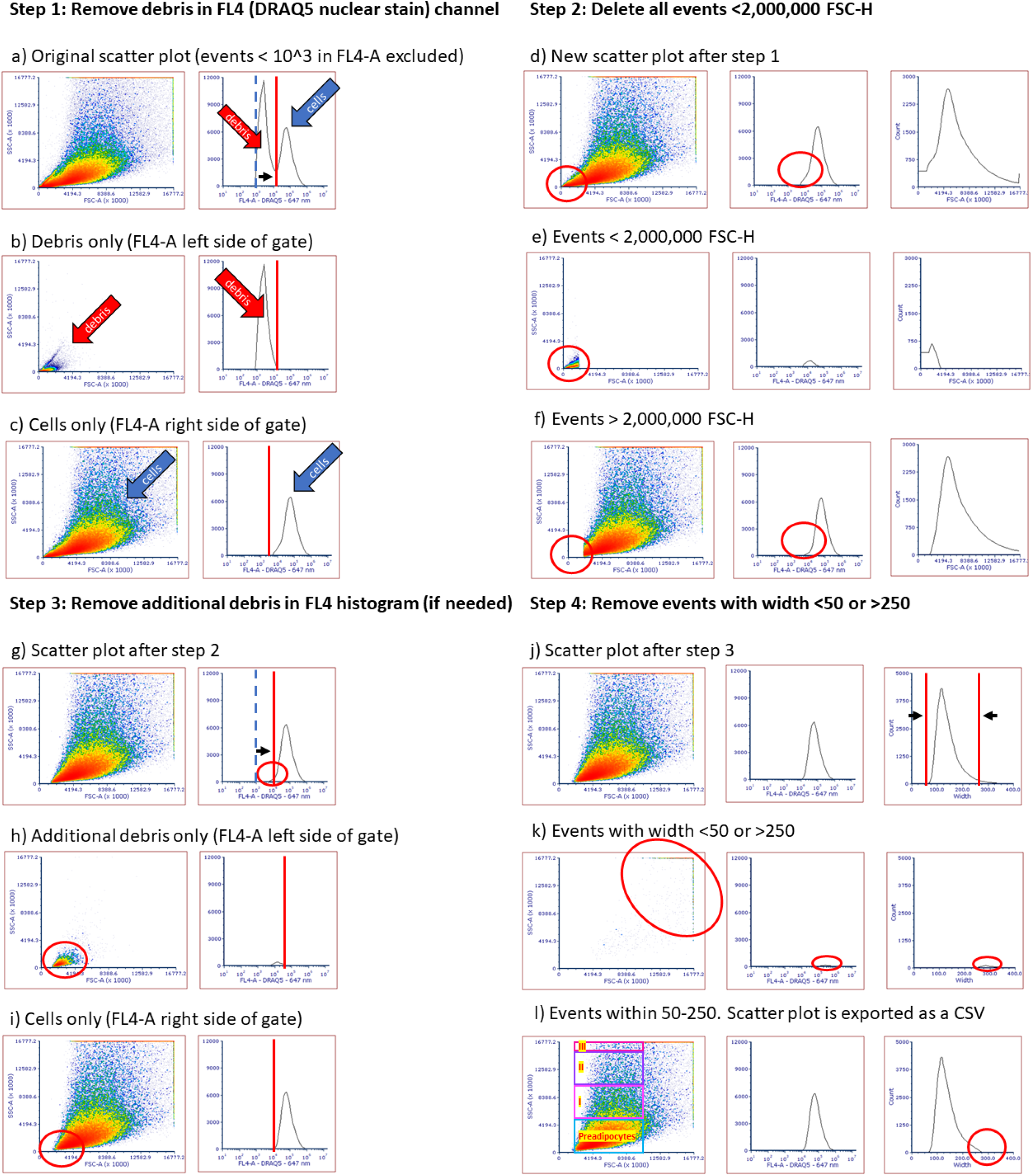
Flow Cytometry Generalized Gating Strategy for Adipocytes. (a) The original scatter plot with events below 10^3 in the FL4-A (647 nm, DRAQ5, nuclear stain) histogram not collected to minimize events from debris (blue dashed line: original gate, red solid line: newly adjusted gate). (b) The left side of the FL4-A histogram gate shows low FSC-A and low SSC-A from debris. (c) The right side of the FL4-A histogram gate shows adipocytes. (d) Newly gated scatter plot after debris is deleted based on the FL4-A histogram gate. (e) Scatter plot shows only events below 2,000,000 FSC-H. (f) Scatter plot with events below 2,000,000 FSC-H removed. (g) The FL4-A histogram gate may need further adjustment if a conservative gate was initially used in Step 1. (h) Additional gating on the left side of the FL4-A histogram gate shows low FSC-A and low SSC-A from debris. (i) Additional gating on the right side of the FL4-A histogram gate shows less debris while displaying adipocytes. (j) Using width histogram to gate events with width less than 50 or greater than 250. (k) Scatter plot showing only events with width less than 50 or greater than 250. (l) Final scatter plot showing only events with width less than 50 or greater than 250. This scatter plot data is exported as a CSV and events are binned into subpopulations (Preadipocytes: 2 million – 10 million FSC-A, 0 – 5 million SSC-A, Group I: 2 million – 10 million FSC-A, 5 million – 10 million SSC-A, Group II: 2 million – 10 million FSC-A, 10 million – 15 million SSC-A, Group III: 2 million – 10 million FSC-A, 15 million – 16.7 million SSC-A).

### Data and Statistical Analysis

Excel (Microsoft) software was used to bin the adipocyte subpopulations based on SSC. The gaiting strategy histograms and scatter plots were generated using FCS Express 7 flow cytometry software (De Novo Software). Prism 8 (Graph Pad) software was used to perform all statistical analyses. All experiments were conducted with at least three biological replicates. The distribution of each data set was analyzed, and the D’Agostino-Pearson test (α=0.05) was performed to test for normality. Comparisons among multiple subpopulations were performed using two-way analysis of variance (ANOVA) or Kruskal-Wallis test with Tukey post hoc testing. All graphs are presented as mean ± standard error of the mean (SEM) unless otherwise stated. Significance was determined according to *p < 0.05, **p ≤ 0.01, ***p ≤ 0.001, and ****p ≤ 0.0001.

## Representative results

### Flow Cytometry Gating Strategy for Adipocytes

Before running samples, the flow cytometer was programmed to discard any events with a signal intensity below 10^3 in the FL4-A (647 nm, DRAQ5, nuclear stain) channel histogram to minimize events from debris. This step was determined experimentally by running samples without any gates applied (data not shown). We observed two distinct peaks in the FL4-A channel histogram, which represent debris (left peak) and cells (right peak) (Fig. 1a). The left peak within the FL4-A channel gate was confirmed to be debris because it has low DRAQ5 fluorescent intensity, low forward scatter (FSC), and low side scatter (SSC) compared to the cells (right peak) (Fig. 1b). The FL4-A channel gate was adjusted to the right to remove the debris (left peak) from the histogram (Fig. 1c). Next, events with <2,000,000 FSC-H were deleted to remove events from debris further (Fig. 1e, f). This FSC-H value was determined experimentally by running adipocytes cultured in 2D (Supplementary Fig. 1). Adipocytes cultured in 2D do not have the same matrix components to be digested; therefore, minimal signal from debris is expected. After removing FSC-H events, the FL4-A channel gate within the histogram may need further adjustment if a conservative gate was initially used (Fig. 1g, h). One distinct peak should now exist within the FL4-A channel histogram, and this is the adipocyte population (Fig. 1i). Next, events with a width >250 or < 50 were removed to avoid adipocyte clumps or doublets (Fig. 1j). This was confirmed experimentally since events with a width above 250 typically showed a greater forward scatter than 10,000,000 (AU) and often saturated both FSC-A and SSC-A detectors (Fig. 1k). The scatter plot data was then exported as a CSV file, and events were binned into subpopulations (Group I: 2 million – 10 million FSC-A, 5 million – 10 million SSC-A, Group II: 2 million – 10 million FSC-A, 10 million – 15 million SSC-A, Group III: 2 million – 10 million FSC-A, 15 million – 16.7 million SSC-A) (Fig. 1l) [23–25]. Mean fluorescence intensity was recorded from the FL1 (533/30) or FL2 channels (585/40).

The subpopulations binning strategy may need to be adjusted based on the flow cytometer and software. Different values can be chosen to better fit scatter plots when binning subpopulations.

### Using Side Scatter (SSC) to Bin Adipocyte Subpopulations

FSC and SSC alone provide useful morphological details about adipocyte population heterogeneity. Theoretically, FSC can be correlated with cell size, while SSC can be correlated with intracellular complexity[26]. To determine if the complexity of intracellular lipid content contributes to SSC, hMSCs underwent adipogenesis for 0-28 days, and flow cytometry was performed (Fig. 2). On day 0 of adipocyte differentiation, virtually all cells (99.99%) were detected in the preadipocyte subpopulation (Fig. 2b-d). After 7 days of adipogenic induction, adipocytes are detected in Groups I-III, and their subpopulation continues to increase through the culture period (Fig. 2b-d). These findings support our hypothesis that there is a correlation between SSC and lipid accumulation since undifferentiated cells only appear in the preadipocyte subpopulation, and the number of adipocytes in Groups I-III increased with time in culture.

**Figure 2.**
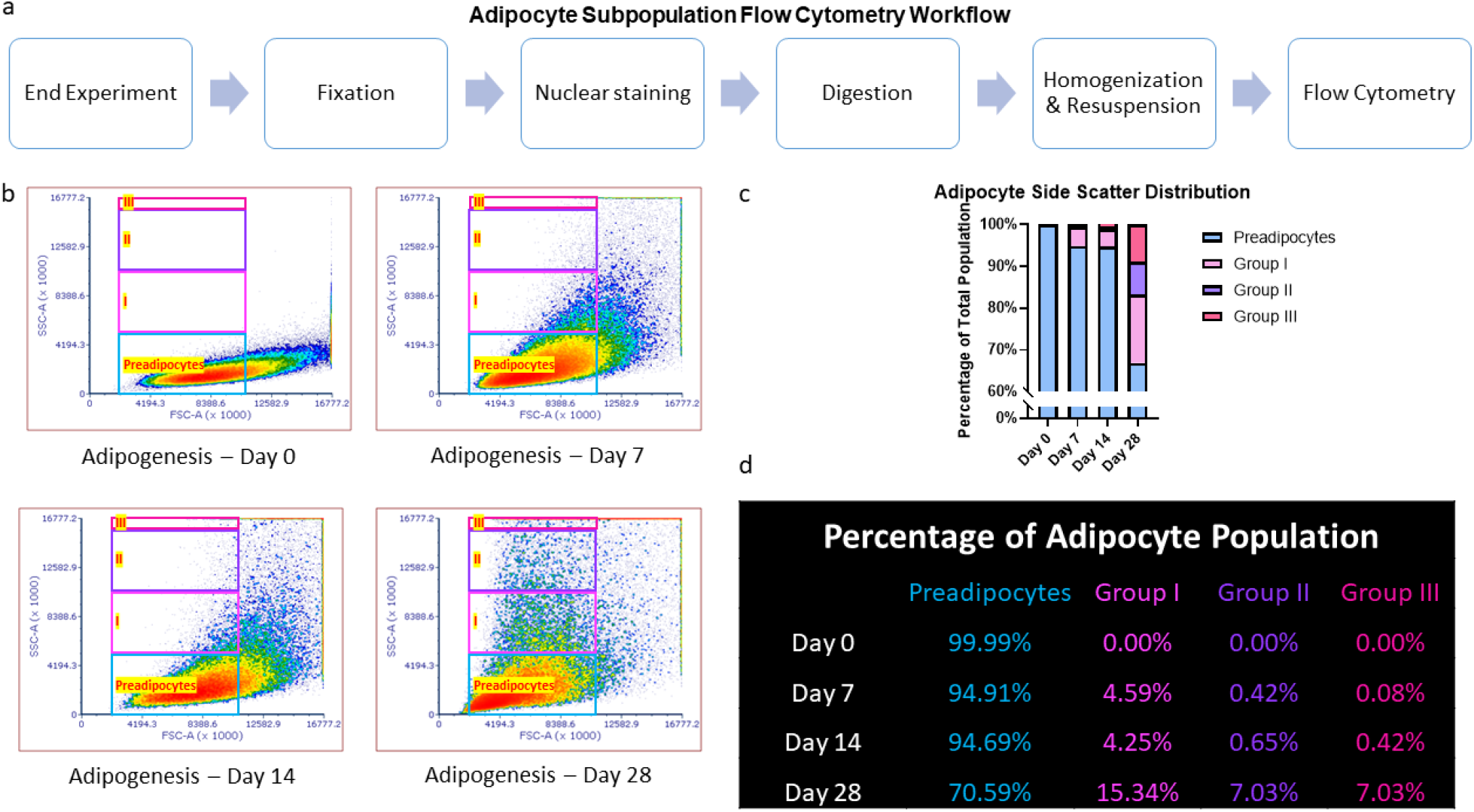
Using Side Scatter (SSC) to Bin Adipocyte Subpopulations. (a) Workflow for running flow cytometry on engineered 3D adipose tissue (AT) constructs for characterizing adipocyte subpopulations. (b) Scatter plots of adipocytes at different time points of adipogenesis (day 0, day 7, day 14, day 28). (c, d) Percentage of each subpopulation in the total adipocyte population. Subpopulations are color coded as follows: Group III – magenta, Group II – purple, Group I – pink, Preadipocyte – light blue.

**Figure 3.**
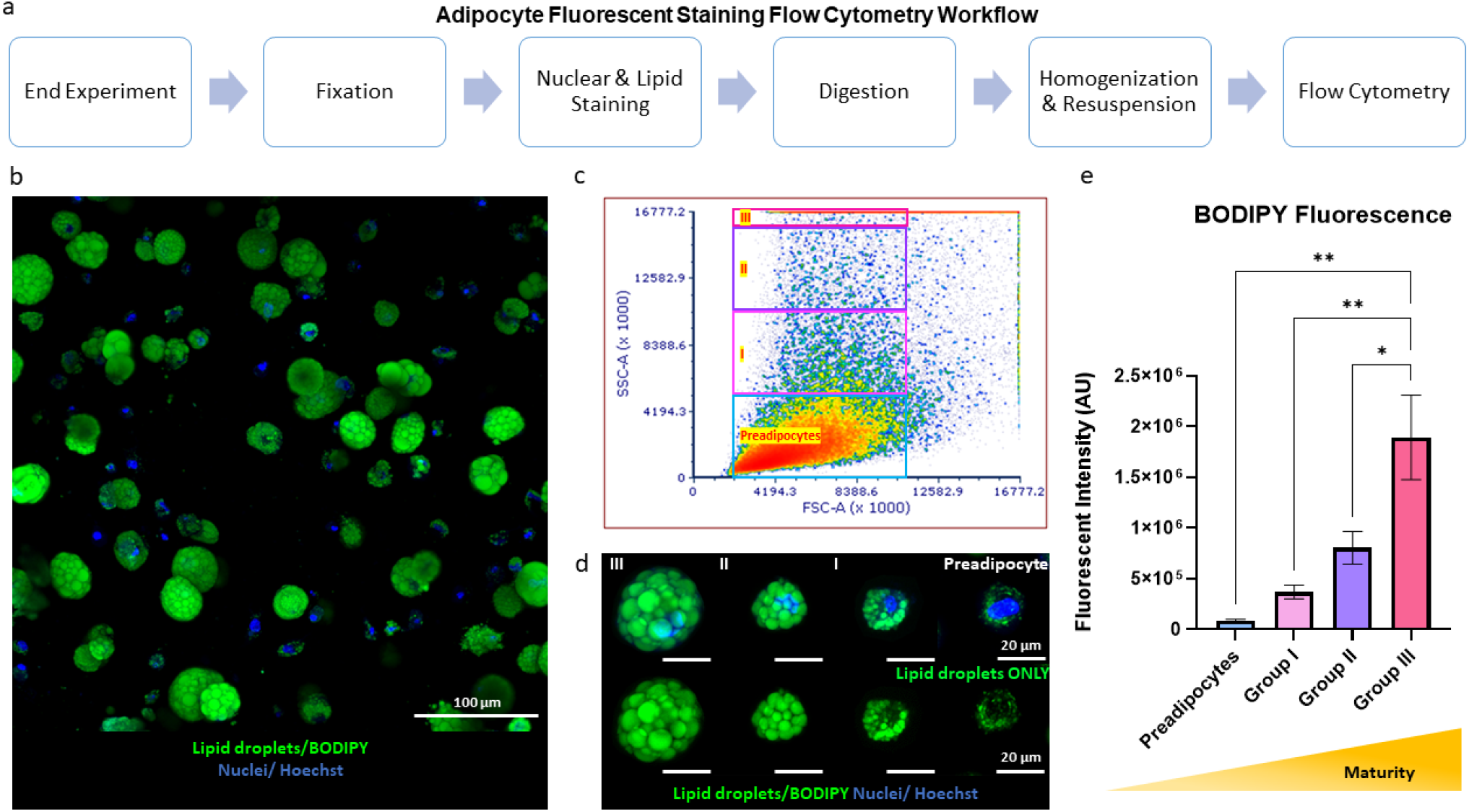
Fluorescent Staining of Adipocytes: BODIPY Fluorescence for Lipid Droplet Content. (a) Workflow for running flow cytometry on 3D AT constructs to quantify lipid droplets. (b) Representative image of adipocyte population within AT construct stained with Hoechst 33342 (nuclei) and BODIPY 493/503 (lipid droplets). (c) Scatter plots of adipocytes after 28 days of culture. (d) Representative images of adipocytes within each subpopulation (Group III, Group II, Group I, Preadipocyte) with nuclei (blue, Hoechst 33342) and lipid droplets (green, BODIPY 493/503) visualized. Top row – overlay of lipid droplets and nuclei. Bottom row – lipid droplets only. (e) Fluorescent intensity of BODIPY in each subpopulation. Subpopulations are color coded as follows: Group III – magenta, Group II – purple, Group I – pink, Preadipocyte – light blue.

**Figure 4.**
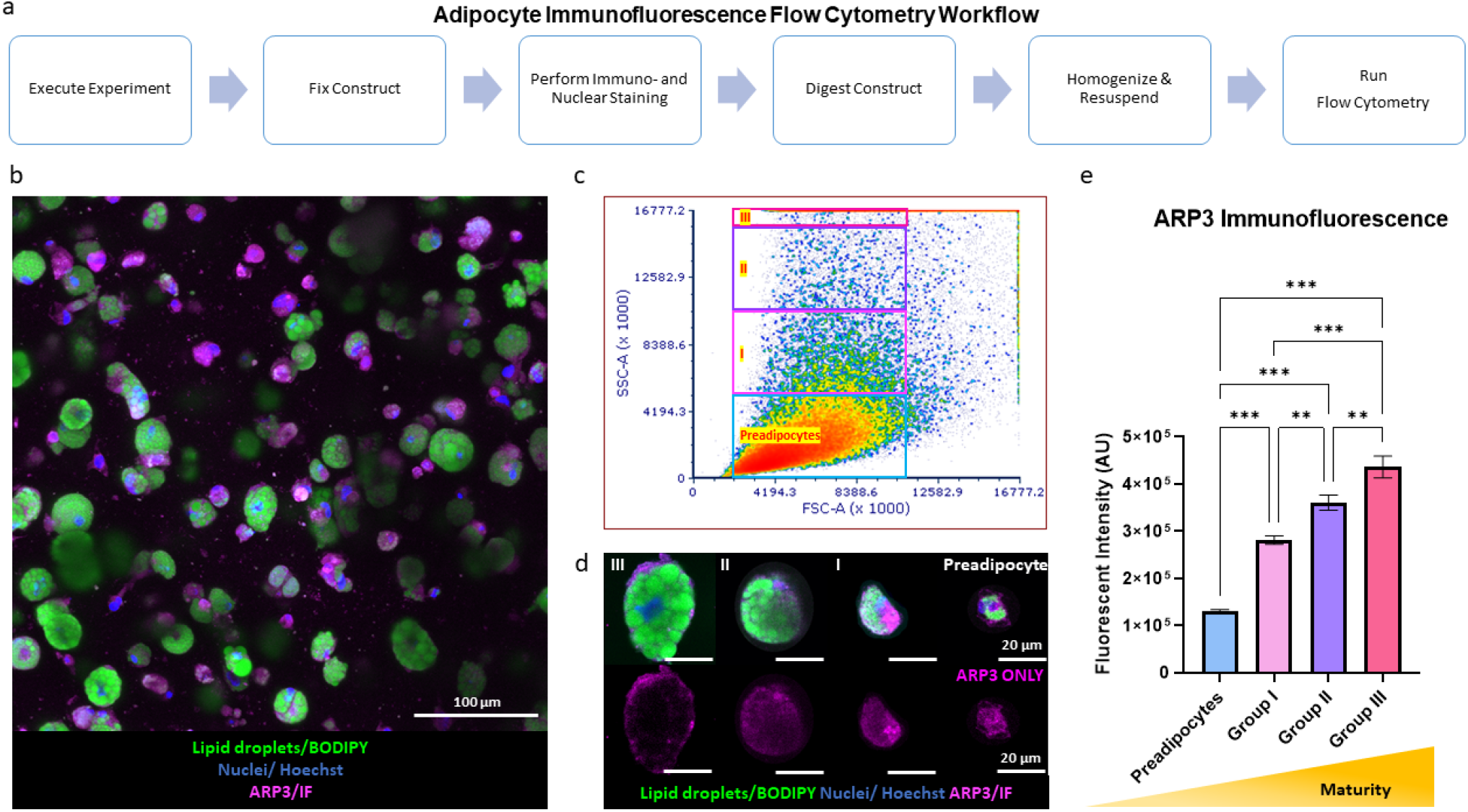
Adipocyte Immunofluorescence: ARP3 Immunofluorescence for Cortical Actin Branching. (a) Workflow for running flow cytometry on 3D AT constructs to quantify adipocyte ARP3 protein content. (b) Representative image of adipocyte population within AT construct with nuclei (blue, Hoechst 33342), lipid droplets (green, BODIPY 493/503), and ARP3 (magenta, immunofluorescence) visualized. (c) Scatter plots of adipocytes after 28 days of culture. (d) Representative images of adipocytes within each subpopulation (Group III, Group II, Group I, Preadipocyte) with nuclei (blue, Hoechst 33342), lipid droplets (green, BODIPY 493/503), and ARP3 (magenta, immunofluorescence) visualized. Top row – overlay of ARP3, lipid droplets, and nuclei. Bottom row – ARP3 only. (e) Fluorescent intensity of ARP3 immunofluorescence in each subpopulation. Subpopulations are color coded as follows: Group III – magenta, Group II – purple, Group I – pink, Preadipocyte – light blue.

**Figure 5.**
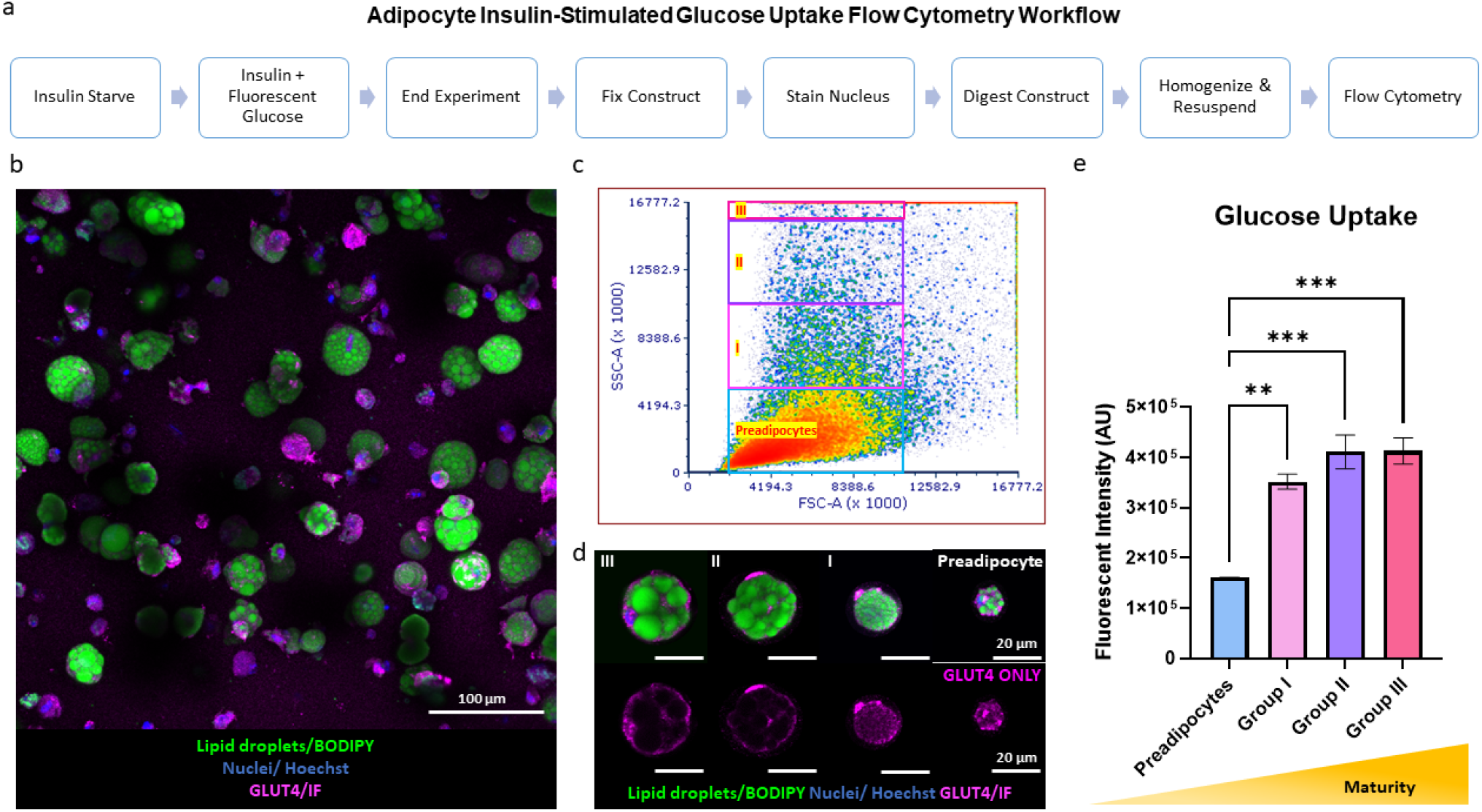
Measuring Adipocyte Function: Insulin-Stimulated Glucose Uptake. (a) Workflow for running flow cytometry on 3D AT constructs for characterizing adipocyte insulin-stimulated glucose uptake. (b) Representative image of adipocyte population within AT construct with nuclei (blue, Hoechst 33342), lipid droplets (green, BODIPY 493/503), and GLUT4 (magenta, immunofluorescence) visualized. (c) Scatter plots of adipocytes after 28 days of culture. (d) Representative images of adipocytes within each subpopulation (Group III, Group II, Group I, Preadipocyte) with nuclei (blue, Hoechst 33342), lipid droplets (green, BODIPY 493/503), and GLUT4 (magenta, immunofluorescence) visualized. Top row – overlay of GLUT4, lipid droplets, and nuclei. Bottom row – GLUT4 only. (e) Fluorescent intensity of 2-NBDG in each subpopulation. Subpopulations are color coded as follows: Group III – magenta, Group II – purple, Group I – pink, Preadipocyte – light blue.

### Fluorescent Staining of Adipocytes: BODIPY 493/503 Fluorescence

Next, the correlation between lipid content and SSC was validated by staining lipids (BODIPY 493/503) of day 28 AT constructs and performing flow cytometry (Fig. 3). BODIPY 493/503 fluorescent intensity was shown to increase linearly with increasing SSC in the adipocyte subpopulations (Fig. 3e). These findings confirm that adipocyte differentiation involves increased intracellular complexity, explicitly derived from adipocyte lipids.

### Immunofluorescence (IF) Staining of Adipocytes: ARP3 Immunofluorescence

Next, we validated that immunofluorescence signal could be quantified with flow cytometry (Fig. 4). ARP3 was visualized by IF, where it should be localized to the cell cortex as adipocytes mature (Fig. 4b, c). ARP3 fluorescent intensity was shown to increase linearly with increasing SSC in the adipocyte subpopulations (Fig. 4e). These findings suggest adipogenesis involves the accumulation of ARP3 dependent cortical actin, playing critical roles in intracellular transport while providing space for intracellular lipid accumulation.

### Measuring Adipocyte Function: Insulin-Stimulated Glucose Uptake

Flow cytometry for adipocyte characterization is not limited to characterizing intracellular features. Flow cytometry can also be used for functional assays, such as insulin-stimulated glucose uptake (Fig. 5). This is advantageous since quantifying insulin-stimulated glucose uptake is not easily discernable in 3D cells in constructs since all cells have some basal uptake. As adipocytes mature, GLUT4 becomes more localized to the cell cortex (Fig. 5d), as it docks onto the increased cortical actin. Performing flow cytometry after insulin-stimulated glucose uptake shows that 2-NBDG fluorescent intensity in adipocytes increases linearly with SSC in the adipocyte subpopulations (Fig. 5e).

### Troubleshooting

## Conclusion

Flow cytometry provides a quick and easy method to analyze heterogeneous adipocyte populations. In this study, we have demonstrated a successful gating strategy for isolating and measuring adipocyte populations. We have shown that adipocyte SSC increases over time as the adipocytes mature, and this was further validated by staining with BODIPY 493/503. Additionally, we have validated that adipocyte maturation is regulated by cortical actin remodeling as branched actin. Lastly, we validated the use of flow cytometry for the functional assay, insulin-stimulated glucose uptake and confirmed that glucose uptake is dependent on cortical actin remodeling. Together, the flow cytometry workflows presented here show the effectiveness of binning individual adipocytes based on their level of maturity and allow for comparisons within subpopulations traditional methods cannot provide.

## Author statement

***Evangelia Bellas:*** *Conceptualization, Methodology, Supervision, Writing – Review & Editing, Project Administration, Funding Acquisition* ***Michael Struss***: *Methodology, Formal Analysis, Validity Tests, Visualization, Investigation, Writing-Original draft preparation*. ***Golnaz Anvari***: *Visualization, Investigation*.

## Acknowledgments

- The authors would like to acknowledge funding support from NASA Space Biology grant 80NSSC19K0427 to E.B.

## Supplementary material *and/or* additional information

**Supplementary Figure 1:**
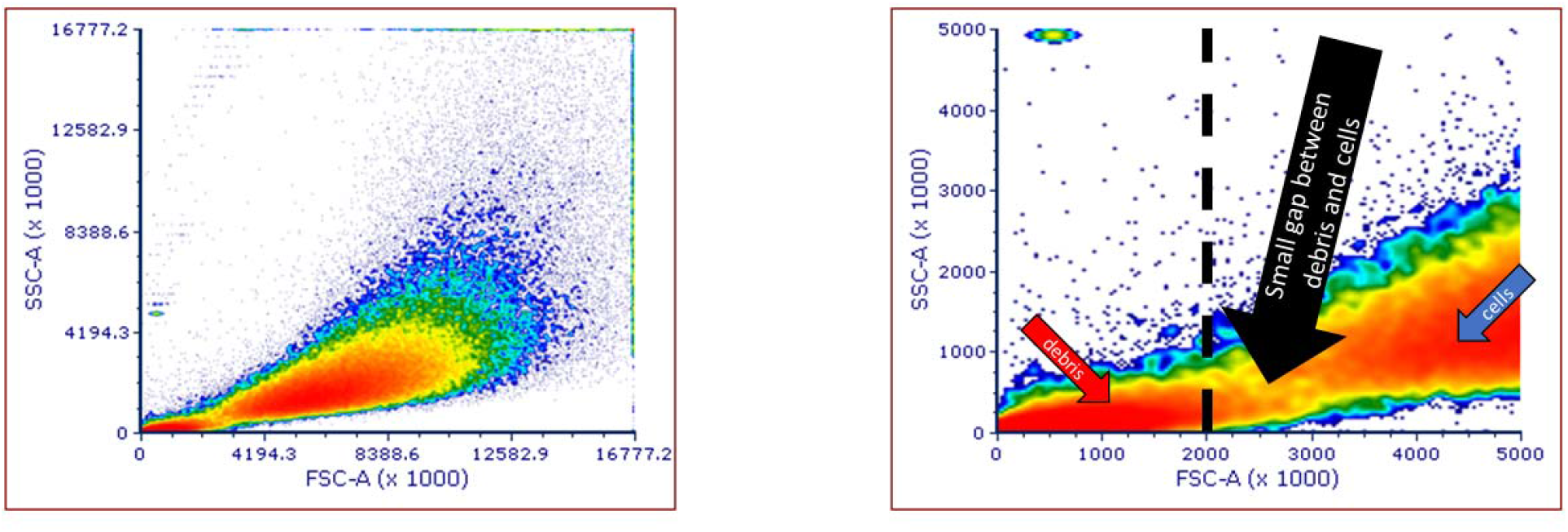
Scatter Plot for Adipocytes Grown in 2D Petri Dishes. Adipocytes were differentiated for 7 days, and flow cytometry was performed. (a) Full scatter plot and (b) zoomed in scatter plot show debris exists below 2,000,000 FSC-A.

